# Differential Inhibition of E-S Potentiation and Long-term Potentiation by Cell-derived and Arctic Amyloid Beta

**DOI:** 10.1101/601492

**Authors:** Adrienne L. Orr, Jason K. Clark, Daniel V. Madison

**Author notes:** Corresponding Author, Beckman Center, B100, 279 Campus Dr., Stanford, California, USA 94305-5345.

## Abstract

Soluble oligomers of amyloid-beta peptide (Abeta) have been implicated in the onset of memory deficits in Alzheimer’s disease, perhaps due to their reported ability to impair long-term potentiation (LTP) of synaptic strength. We previously showed the effect of Abeta on LTP depends on the strength of LTP induction. Furthermore, Abeta affects EPSP-Spike (E-S) potentiation more robustly than LTP, suggesting that E-S potentiation may be equally important to learning and memory in the context of Alzheimer’s disease. Here we extend our findings to two additional forms of Abeta that form higher concentrations of soluble Abeta oligomers and show that they also affect E-S potentiation at induction strengths where there is no effect on LTP in hippocampal slices.

## Introduction

Alzheimer’s disease (AD) is a progressive neurodegenerative disease characterized by memory impairments and cognitive decline. It is widely accepted that soluble oligomers of amyloid-beta peptide (Abeta) may mediate the memory deficits observed in early Alzheimer’s disease [1, 2]. In particular, Abeta has been shown to disrupt long-term potentiation (LTP) of synaptic strength *in vivo* and *in vitro* [3-7].

However, we recently showed that Abeta affects EPSP-Spike (E-S) potentiation much more strongly than it does LTP [8]. E-S potentiation is a form of intrinsic plasticity induced concurrently with synaptic LTP [9]. It is believed to be expressed as a change in the electrical coupling between the dendritic synaptic inputs and the soma, such that a greater proportion of the excitatory post-synaptic potential (EPSP) survives at the spike trigger zone [10]. Because E-S potentiation generates a greater action potential output for a given synaptic EPSP, it provides an additional boost to the efficacy of the EPSP beyond the potentiation (LTP) that occurs at the synapse. Interestingly, though E-S Potentiation has received far less attention than its sibling LTP, it was described alongside LTP in the earliest reports of this plasticity [9]. Although it is induced concurrently with synaptic LTP, it is thought to be a mechanistically distinct process. Nevertheless, some induction features are shared, and the two may share importance in the mechanisms underlying memory storage.

Interestingly, we have confirmed that LTP is reduced in the presence of Abeta after a weak induction protocol (five trains of four stimuli at 100 Hz, 400 ms inter-train interval), but is impervious to Abeta after a relatively strong induction protocol (three trains at 100 Hz for 1 s, 15 ms inter-train interval) [8]. In contrast, E-S potentiation was reduced in the presence of Abeta, under all conditions tested, regardless of the strength of the induction protocol.

Previously, we showed that synthetic Abeta1-42 peptide inhibits E-S potentiation (and weakly-induced LTP) indirectly by blocking endocannabinoid-mediated disinhibition [8]. In this paper, we extend these findings to two additional forms of Abeta that form high concentrations of soluble oligomers: a mutant, cell-derived amyloid beta, and synthetic Arctic Abeta1-40, by measuring the effect of sub-toxic concentrations of all three forms of Abeta on E-S potentiation.

## Materials and Methods

### Slice Preparation

All experiments were performed using male Wistar rats (P25-P40, Harlan Laboratories, Indianapolis, IN). Animals were anesthetized with halothane or isoflurane (Sigma) and swiftly decapitated in accordance with institutional regulations. Brains were rapidly removed and the hippocampus was dissected out in ice-cold oxygenated (95% O_2_-5% CO_2_) high-sucrose/low-sodium slicing solution composed of (in mM): 2.5 KCl, 6.0 MgSO_4_, 2.5 CaCl_2_, 1.0 NaH_2_PO_4_, 26.2 NaHCO_3_, 11.0 glucose, and 210.0 sucrose (Sigma). Transverse hippocampal slices were cut to 400-µm using a manual tissue chopper (Stoelting Co, Wood Dale, IN) moistened with chilled slicing solution. Slices recovered in a custom-made submersion chamber filled with artificial cerebral spinal fluid (ACSF) composed of (in mM): 119 NaCl, 2.5 KCl, 6.0 MgSO_4_, 2.5 CaCl_2_, 1.0 NaH_2_PO_4_, 26.2 NaHCO_3_, and 11.0 glucose (Sigma). The ACSF was pre-saturated by 95% O_2_/5% CO_2_ at room temperature. After equilibrating for at least 1.5 hours, slices were transferred to a submerged recording chamber and continuously superfused with 95% O_2_/5% CO_2_ saturated ACSF (1.5 – 2 mls/min) at room temperature. All peptides were bath applied in oxygenated ACSF.

### Electrophysiology

To elicit field potentials, the Schaffer collateral/commissural afferent (SC/AF) pathway was stimulated using a concentric bipolar platinum-iridium electrode (Frederick Haer & Co, Bowdoinham, ME) placed in the stratum radiatum of the CA1 region. Glass microelectrodes were placed in stratum radiatum or stratum pyramidal 1-2 millimeters away from the stimulating electrode. The microelectrodes were pulled from borosilicate glass (1.65 mm O.D. − 1.15 mm I.D.) using a Flaming-Brown horizontal microelectrode puller (Model P-87, Sutter Instruments, Novato, CA) and were filled with 3M NaCl, giving an input resistance of 1-5MΩ. The baseline stimulus intensity was set so that the population spike area was 10% of maximum. fEPSPs were elicited by delivering 0.1 ms pulses at 0.033 Hz (every 30 s) to the SC/AF fibers. A stable baseline fEPSP was achieved for at least 20 minutes before the start of each experiment. Slices with variable or drifting baselines were discarded. For the induction of LTP and E-S potentiation, high-frequency stimulation (HFS) consisting of three trains of 100 Hz for 1 s with an inter-train interval of 15 seconds was delivered at baseline stimulus intensity.

### Data Acquisition and Analysis

Analog signals were low-pass filtered at 3 kHz, and digitized at a rate of 10 kHz (Axoclamp 2-A amplifier, Brownlee Precision model 200 amplifier, National Instruments data acquisition board, LabVIEW software). All data were analyzed offline using a custom-written LabVIEW based program. LTP and E-S potentiation were calculated by normalizing each data point to the 10 min immediately before HFS, and averaging the data between 30-40 minutes post-HFS. E-S coupling was calculated from the ratio of the area of the population spike to the initial slope of the fEPSP. E-S potentiation is represented by the change in E-S coupling in response to a tetanizing stimulus. The increase observed in the population spike in relation to the EPSP slope represents E-S potentiation because decreasing the stimulus to reduce the EPSP field back to its baseline size still produced a population spike greater than control (not shown).

Statistical data are reported as means ± standard error of the mean (S.E.M.). Statistical significance was determined by one- or two-tailed Student’s T-tests, and significance level was set at *P* < 0.05 or 0.01 (marked by * or **, respectively).

### Cell-derived and synthetic Abeta peptides

Cell-derived Abeta was prepared as described previously [2, 4] from 7PA2-cell conditioned media (CM), a CHO cell line that stably expresses human APP751 with the V717F mutation. Control experiments used CM from un-transfected CHO cells. Briefly, cells were grown to confluence in Dulbecco’s modified Eagle’s medium (DMEM) supplemented with 10% fetal bovine serum, penicillin/streptomycin, L-glutamine, and G418. Once confluent, cells were washed and cultured overnight in serum-free DMEM. The media was collected and centrifuged at 1000 g with protease inhibitors. When sufficient CM was collected, it was centrifuged (3000 g) at 4C in YM-3 Centricon tubes to concentrate proteins larger than 3 kDa. The concentrated CM was thawed and diluted 1:15 in ACSF immediately before use.

Stock solutions of synthetic Abeta1-42 (50 μM in 0.1% NH_4_OH) and Arctic Abeta1-40 (1 mg/ml in water) were diluted to 500 nM in oxygenated ACSF. Vehicle control was prepared from a stock solution of 0.1% NH_4_OH. For all experiments, Abeta or control solution was applied for 40 minutes beginning 20 minutes prior to HFS. In experiments using CM or 7PA2, application began 30 minutes prior to HFS. All peptide stocks were stored at –80 C, and –20 C several days before use.

## Results

In our previous work, we showed that the mechanism by which Abeta1-42 impairs both LTP and E-S potentiation is indirect. The direct action of Abeta1-42, we previously discovered, is to block endocannabinoid-mediated disinhibition [8]. This preserves synaptic inhibition during high-frequency stimulation that is used to induce both LTP and E-S potentiation, blunting the postsynaptic voltage summation that is necessary to induce LTP, opposing its induction. Thus, Abeta1-42 influences whether summated EPSPs can reach the threshold for induction of LTP without inhibiting LTP directly. E-S potentiation on the other hand, while also blocked by Abeta1-42, does not appear to have the same induction threshold characteristic as LTP. In an effort to better understand how Abeta may be influencing E-S potentiation, we wanted to determine if variants of Abeta with different properties would also impair E-S potentiation similar to wild-type Abeta1-42, under conditions where LTP is spared. In our current work, we measured extracellular field potential recordings in acute hippocampal slices in the presence of two different Abeta peptide variants known to form higher concentrations of oligomers and protofibrils than what is typically found with synthetic Abeta1-42, to determine their effects on the induction and maintenance of E-S potentiation and LTP.

We first examined Abeta generated from the 7PA2 CHO cell line. This cell line expresses human APP with a V717F mutation, resulting in the production of a higher than normal concentration of stable, low-N Abeta oligomers [3]. As a result, 7PA2 has been reported to be a more-potent disruptor of LTP than synthetic Abeta1-42 [4]. Hippocampal slices were superfused with either 1X 7PA2- or CHO-conditioned medium not containing 7PA2 for 30 minutes prior to HFS. We found that LTP in 7PA2 treated slices did not differ from those slices treated with CHO only (Figure 1A, Figure 3). However, the ratio of population spike to EPSP slope showed that 7PA2 conditioned media impairs E-S potentiation in the same manner as synthetic Abeta (Figure 1C and Figure 4).

**Figure 1:**
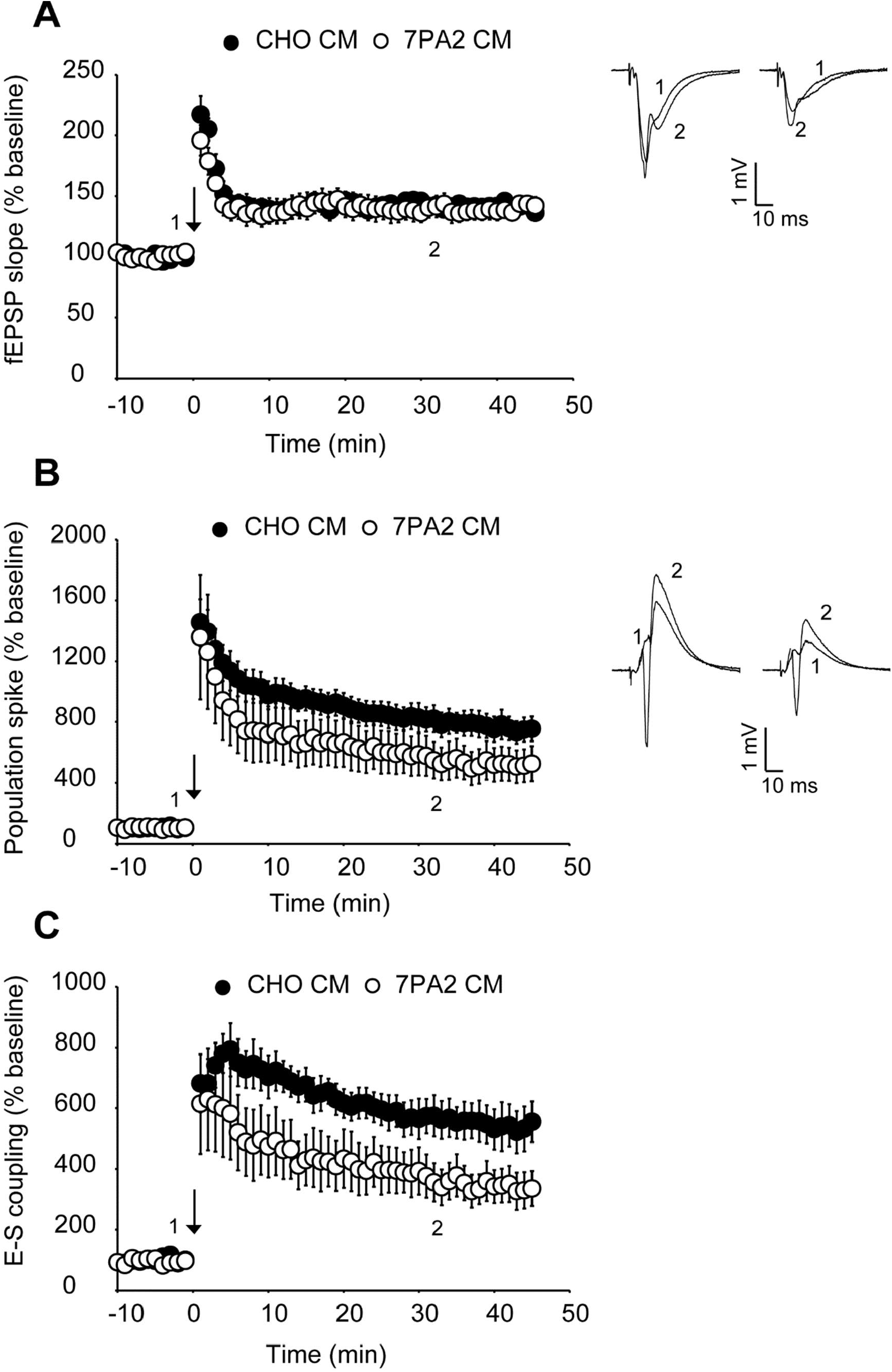
Cell derived Abeta impairs E-S potentiation but not synaptic potentiation. Cell-derived Abeta (1X conditioned media from 7PA2 cell-line, which is CHO transfected with hAPP V717F mutation) (○; n = 5) untransfected CHO-conditioned media (•; n = 6) was superfused over slices while fEPSPs were recorded in the stratum radiatum and stratum pyramidal of the CA1 region. After 30 minutes of perfusion, high-frequency stimulation (HFS, ↓) was delivered to induce plasticity. HFS consisted of 3 1s 100 Hz trains with 15 s intervals between trains. Data are normalized to the 10 minutes before HFS and expressed as means ± standard error of the mean. A) Cell-derived Abeta does not impair potentiation of fEPSP slope (•; 142 ± 4, ○; 139 ± 8, p = 0.365). B) Population spike potentiation is not significantly changed by cell-derived Abeta (•; 791 ± 82, ○; 536 ± 118, p = 0.055). C) E-S coupling potentiation is impaired by cell-derived Abeta (•; 558 ± 62, ○; 353 ± 61, p = 0.021). Right of A and B: Example EPSPs from CHO CM (left) and cell-derived Abeta (right) treated slices.

We next looked at an alternative Abeta formulation, the Artic Abeta1-40. The Arctic mutation (G22) of amyloid precursor protein causes aggressive familial neuropathology associated with Alzheimer’s-like dementia due to enhanced protofibril formation [11]. Arctic Abeta1-40 is a peptide formulation of Abeta that has also been shown to be more potent in impairing LTP than wild-type Abeta1-42 [12] *in vivo*. As with 7PA2 cell-derived Abeta, we found Arctic Abeta1-40 also impaired E-S potentiation but did not impair LTP with strong induction (Figures 2, 3 and Figure 4). Overall, when compared to the vehicle controls, all three Abeta formulations tested robustly and reliably impaired E-S potentiation (Figure 4). Conversely, in no case did we detect a reduction in LTP (Figure 3), regardless of the fact that these peptide formulations were clearly active against E-S potentiation.

**Figure 2:**
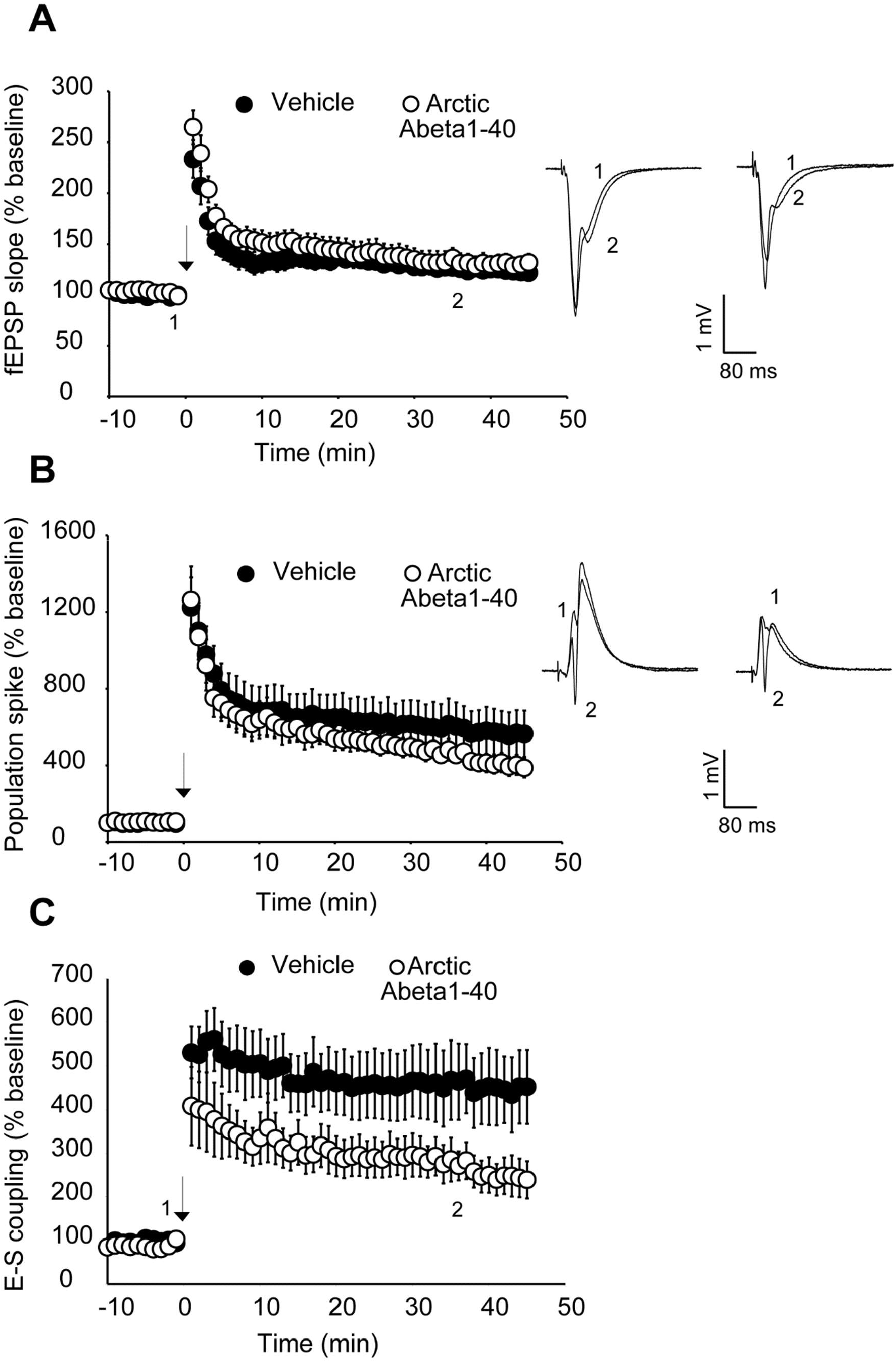
Arctic Abeta1-40 impairs E-S potentiation, but not synaptic potentiation. 500 nM Arctic Abeta1-40 (Abeta1-40 with E22G mutation) (○; n = 6) or vehicle (•; n = 6) was superfused over slices while fEPSPs were recorded in the stratum radiatum and stratum pyramidal of the CA1 region. After 20 minutes of perfusion, high-frequency stimulation (HFS, ↓) was delivered to induce plasticity. HFS consisted of 3 1s 100 Hz trains with 15 s intervals between trains. Data are normalized to the 10 minutes before HFS and expressed as means ± standard error of the mean. A) Arctic Abeta1-40 does not impair potentiation of fEPSP slope (•; 129 ± 6, ○; 136 ± 11, p = 0.418). B) Population spike potentiation is impaired by Arctic Abeta1-40 (•; 678 ± 109, ○; 458 ± 50, p = 0.033). C) E-S coupling potentiation is impaired by Arctic Abeta1-40 (•; 519 ± 71, ○; 306 ± 45, p = 0.008). Right of A and B: Example EPSPs from vehicle (left) and Arctic Abeta1-40 (right) treated slices.

**Figure 3:**
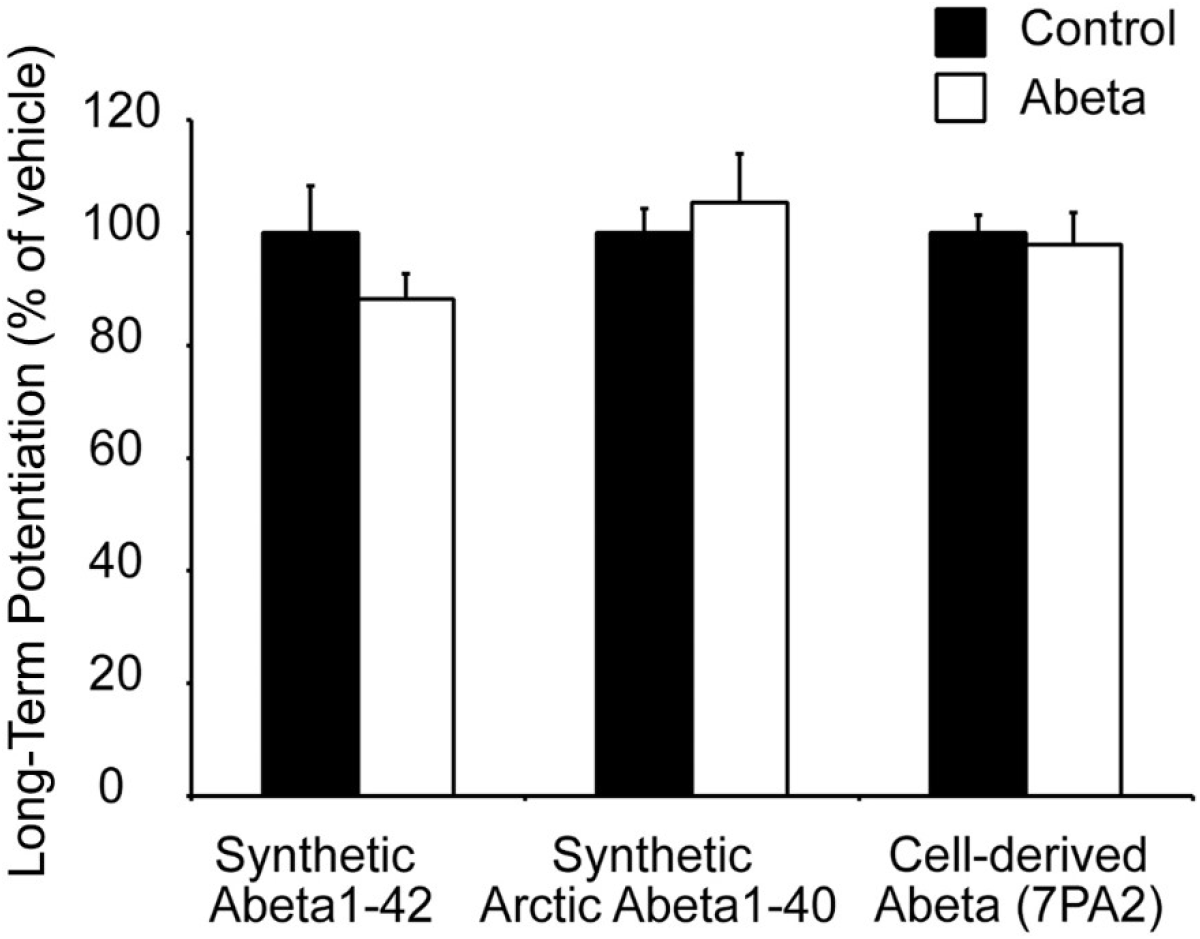
Abeta peptides do not impair LTP. Bar representation of Long-term potentiation with three formulations of Abeta (white bars). Abeta peptides ((500 nM synthetic Abeta1-42 (n = 7), 500 nM synthetic Arctic Abeta1-40 (E22G mutation) (n = 5), and 1X cell-derived Abeta in conditioned media (7PA2 cell line, human APP V717F mutation) (n = 5) (white bars)) have no effect on LTP relative to control ((0.1% NH_4_OH (n = 6), water (n = 6), CHO conditioned media (n = 5) (black bars)) at 30-40 minutes following HFS (p = 0.113, 0.418, 0.365). Data are normalized to the vehicle controls and expressed as means ± standard error of the mean.

**Figure 4:**
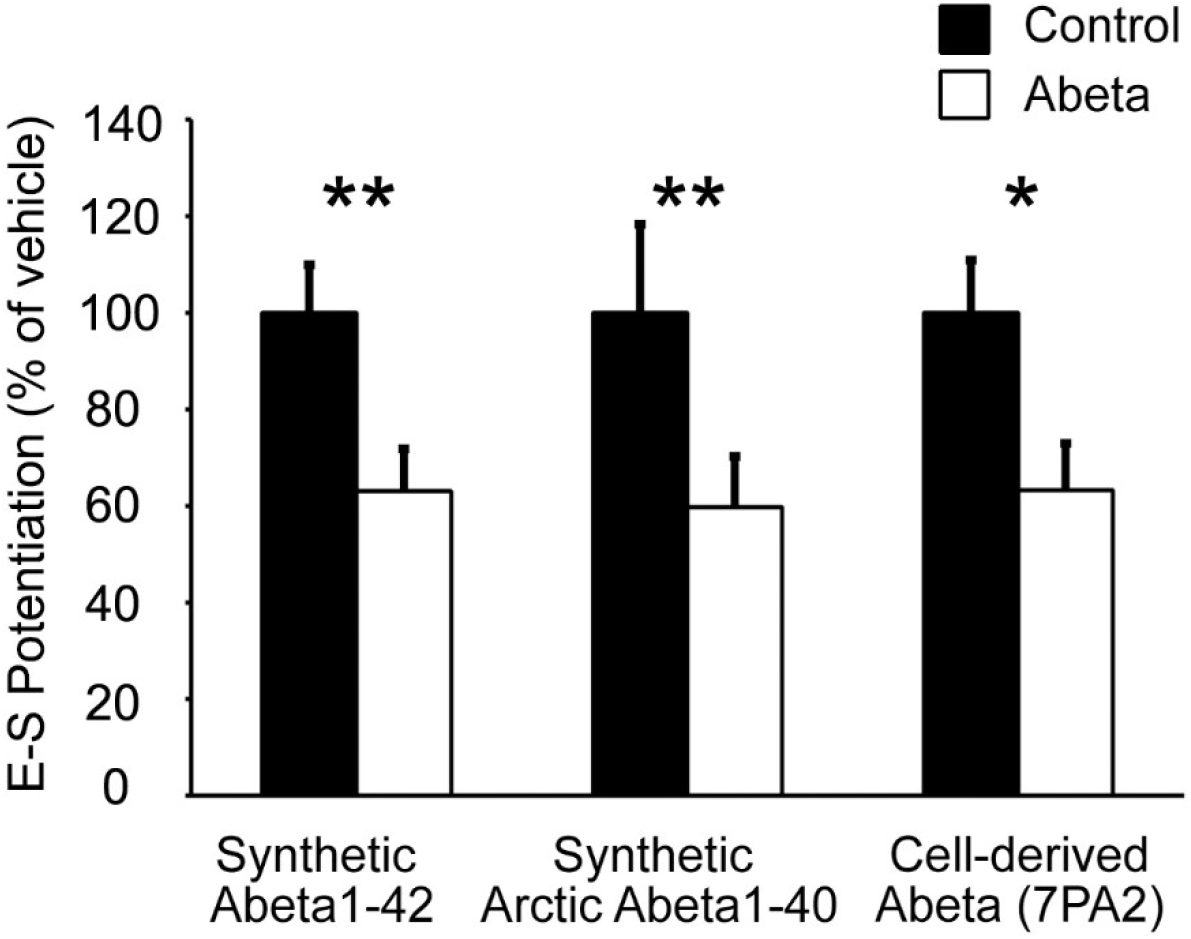
E-S coupling potentiation is robustly impaired by Abeta peptides. Bar representation of E-S potentiation with three different formulations of Abeta (white bars). Abeta peptides ((500 nM synthetic Abeta1-42 (n = 7), 500 nM synthetic Arctic Abeta1-40 (E22G mutation) (n = 5), and 1X cell-derived Abeta in conditioned media (7PA2 cell line, human APP V717F mutation) (n = 5) (white bars)) potently impair E-S potentiation compared to control ((0.1% NH_4_OH (n = 6), water (n = 6), CHO conditioned media (n = 5) (black bars)) at 30-40 minutes following HFS (p = 0.007, 0.007, 0.021). Data are normalized to the vehicle controls and expressed as means ± standard error of the mean.

## Discussion

Memory impairment is one of the hallmark characteristics of Alzheimer’s disease, and early memory deficits are likely due to neuronal dysfunction in the hippocampus, rather than cell death. It is believed that soluble oligomers of Abeta peptide, a cleavage fragment from the amyloid precursor protein, may mediate this dysfunction by impairing a variety of neuronal functions, including synaptic plasticity [13]. Multiple studies have shown that Abeta impairs LTP [3-7, 14], and studies from our own lab have shown exogenously applied Abeta1-42 also impairs E-S Potentiation, and does so with greater potency [8]. While it is known that Abeta1-42 can impair LTP and E-S potentiation, we wished to determine if two different Abeta variants also impair LTP and E-S potentiation similarly to wild-type Abeta1-42.

LTP and E-S potentiation have distinctly different molecular mechanisms, but share common induction features. Here we demonstrate that two additional Abeta variants impair E-S potentiation via a molecular pathway independent of LTP since LTP is spared under our induction protocol. Our data did not replicate previous findings by others that Abeta impairs LTP, even using formulations that are known to have higher oligomer content. We theorize that this discrepancy may be due to the nature of plasticity impairment by Abeta. In a previous work, we demonstrated Abeta inhibits suppression of inhibition through a cannabinoid dependent signaling pathway [8]. In suppressing disinhibition, Abeta acts as a negative modulator of plasticity, interacting with the ability of summating EPSPs to reach the LTP threshold, rather than directly inhibiting the LTP mechanism. As was the case with previously published data using synthetic Abeta1-42, these obstensively more potent forms of Abeta also showed E-S potentiation was not spared, despite the strong induction protocol used. This suggests that Abeta blocks E-S potentiation by a different, more potent mechanism, than the anti-disinhibitory mechanism by which it impairs LTP. These results also suggest that a blockade of E-S potentiation may be a more important factor in Alzheimer’s-associated memory impairment than a reduction in LTP.

In summary, we show that high-oligomer Abeta peptide formulations robustly impair E-S potentiation, but not synaptic potentiation, similar to wild-type Abeta1-42. The ability to properly regulate the coupling of synaptic inputs to the generation of neuronal output (spikes) is essential to normal neural network function. This type of coupling and plasticity may potentially be as, if not more, important as synaptic plasticity for learning and memory, and impairments in E-S potentiation may explain some of the early deficits observed in Alzheimer’s disease. Further study is needed to better understand the effects of Abeta on different types of plasticity other than synaptic plasticity, in order to better understand the nature of Alzheimer’s disease progression, and provide insight for future therapeutic strategies.

## Acknowledgments

This work was supported by an Innovation Grant from Elan Pharmaceuticals, grants from the National Institute of Mental Health (MH065541 and MH111768) and by the Harold and Leila Y. Mathers Charitable Foundation. We thank Sarah Wright, Peter Seubert, Kelly Johnson-Wood and the late Dale Schenk for their valuable assistance with this project. In remembrance of Dale B. Schenk.

